# Structure and its transformation of elliptical nege-like virus Tanay virus

**DOI:** 10.1101/2023.01.20.523382

**Authors:** Kenta Okamoto, Chihong Song, Miako Sakaguchi, Christina Chalkiadaki, Naoyuki Miyazaki, Takeshi Nabeshima, Kouichi Morita, Shingo Inoue, Kazuyoshi Murata

## Abstract

Negeviruses that infect insects are recently identified virus species that are phylogenetically related to several plant viruses. They exhibit a unique virion structure, an elliptical core with a short projection. Negeviruses encode two structural proteins, a glycoprotein that forms a short projection, and an envelope protein that forms an elliptical core. The glycoprotein has been reported only in the negeviruses’ genes, and not in phylogenetically related plant viruses’ genes. In this report, we first describe the three-dimensional electron cryo-microscopy (cryo-EM) structure of Tanay virus (TANAV), one of the nege-like viruses. TANAV particle demonstrates a periodical envelope structure consisting of three layers surrounding the centered viral RNA. The elliptical core dynamically changes its shape under acidic and even low detergent conditions to form bullet-like or tubular shapes. The further cryo-EM studies on these transformed TANAV particles reveal their overall structural rearrangement. These findings suggest putative geometries of TANAV and its transformation in the life cycle, and the potential importance of the short projection for enabling cell entry to the insect hosts.

**Impact statement:** Negeviridae has recently been declared as a virus family that includes virus species exhibiting a unique particle structure that differs from other known viruses. They are known to be common mosquito viruses isolated around the world, but also phylogenetically related to several plant viruses that impair crop production. Therefore, the negeviruses may also play a role in plant ecosystems that threaten agriculture. However, the mechanism of infection and assembly of the negeviruses as well as their structure were unknown. In this study, intact and dissociated structures of the TANAV were first examined using cryo-EM single particle analysis (SPA) and electron cryo-tomography (cryo-ET). These results reveal new structural geometries of the TANAV particle and its dynamic transformation under acidic and even low-detergent conditions, providing new insights into the infection and assembly mechanism in negeviruses.

## Introduction

Plants and their transmission vectors have been interrelated for over 450 million years. Viruses as transmission vectors are an important element of plant ecosystems, but they also cause serious damage to the crop production industry (1). Insects are also higher-level vectors that transmit plant viruses between plants by ingesting their extracts (2). Many plant viruses have been reported to be transmitted by various insects such as leafhoppers, planthoppers, aphids, whiteflies, and thrips (2), but it is unclear whether mosquitoes can also transmit such viruses between plants.

Unclassified taxa of positive single-stranded ((+)ss) RNA negeviruses, including Tanay virus (TANAV), are widely distributed across mosquito species (3–13). Two genus-level taxon of Sandewavirus and Nelorpivirus are suggested in the negeviruses because of the sequence difference in their RdRp gene (7). Negeviruses are also phylogenetically related to crop pathogens such as the Citrus leprosis virus C (CiLV-C), and Blueberry necrotic ring blotch virus (BNRBV) (3, 14). Since the associations between the mosquito negeviruses and the plant viruses have not yet been elucidated, it remains unclear whether the negeviruses are exclusively maintained in mosquitoes or transmitted between mosquitoes and plant hosts (3, 14). The TANAV genome encodes three putative open reading frames (ORFs) (3). ORF1 encodes non-structural proteins for viral replication such as RNA-dependent RNA polymerase (RdRp) (3, 12). ORF2 encodes a protein that is composed of 592 amino acids; its homologs in other nege-like viruses encode structural glycoproteins (3, 8, 12, 13). ORF3 encodes a putative membrane protein of the enveloped core with a predicted molecular weight of approximately 25 kDa (3, 8, 13). Some plant viruses, such as virga- or nege-like viruses, have an enveloped core with glycoprotein spikes that may facilitate the entry of the virus into insect cells; in contrast, other phylogenetically related plant-specific (+)ssRNA viruses do not contain these putative glycoproteins in the genome (14–16). Structural analysis of TANAV particles helps to understand the ability of negeviruses to propagate in mosquito hosts. TANAV particles that are revealed by negative stain EM demonstrated an elliptical shape unlike known mosquito virus particles (3). Later, a unique elliptical core was found in a phylogenetically closely related Castlerea virus (CsV) (13).

Here, we first describe the intact structure of negevirus and its transformation that is caused by acidic and even low detergent buffer conditions. We present the reconstructed three-dimensional (3D) structure of the elliptical core of the TANAV particle and the transformed bullet-like and tubular particles by cryo-EM SPA or cryo-ET at nearly native conditions. These findings suggest a novel structure of the TANAV elliptical core including its transformation, and the putative function of the short projection that is extended from the elliptical core.

## Methods

### Cell culture and sample purification

C6/36 mosquito cells were cultured in minimum essential medium (MEM) supplemented with 10% fetal bovine serum (FBS) with penicillin and streptomycin. The TANAV was inoculated by micro bead attachment to C6/36 cells upon stirring with a spinner flask. After three days of inoculation, the infected culture fluid (ICF) was centrifuged at 13,000 x*g* at 4°C for 30 min. The 1.8 L of ICF was then mixed with 108 g of PEG6000 and 40 g of NaCl and incubated overnight at 4°C. The next day, the ICF was centrifuged at 13,000 ×*g* at 4°C for 30 min, and then the supernatant was removed. The pellet was resuspended in STE buffer (10 mM Tris-HCl, 100 mM NaCl, 1 mM EDTA, pH 7.4). A 15–50% (w/v) sucrose gradient was applied to the suspended sample and ultracentrifuged at 60,000 ×*g* at 4°C for 14 hr. The sample was then fractionated by monitoring protein concentrations via absorbance at 280 nm. The virus fractions were pooled and diluted with STE buffer and ultracentrifuged at 17,000 ×*g* at 4°C for 3 hr. Then, TANAV fractions were dialyzed in 1 L of STE buffer (pH 8), and purified particles were visualized and confirmed using negative staining and transmission electron microscopy. The ORF3 membrane protein band is clearly visible in a purified TANAV fraction, whereas the ORF2 glycoprotein band is difficult to detect due to the high background of cellular proteins carried over during the purification and due to the small abundance (Supplementary Fig. S1A). However, the ORF2 glycoprotein band is visible in a lectin blotting after the sucrose gradient purification (Supplementary Fig. S1B). The purified TANAV fractions displaying both ORF2 and ORF3 protein bands were used for acquiring raw particle images by cryo-EM.

### Cryo-EM data acquisition and single particle analysis (SPA)

Purified TANAV particles were loaded onto a R1.2/1.3 Mo Quantifoil holey carbon grid (Quantifoil Micro Tools) and flash-frozen in liquid ethane using a Vitorobot Mark IV (Thermo Fisher Scientific). The frozen grid was observed with a 200 kV electron microscope (JEM-2200FS, JEOL, Inc.) equipped with a DE-20 direct detector CMOS camera (Direct Electron LP). Omega-type energy filter with a slit width of 20 eV was used for data collection by zero-loss imaging, and TANAV particles images were viewed at a nominal magnification of 30,000 × (1.992 Å/pixel on specimen) and a total electron dose of 20 e-/Å^2^. A total of 75 movie frames were collected for each image. Collected images not including the initial 3 frames were merged after motion correction by a script provided by manufacturer. The contrast transfer function (CTF) was then determined on the merged images by CTFFIND4 (17). After CTF estimation, 19,242 particles were manually selected from the images using RELION 2.1 (18). These particles were classified in two-dimensions (2D) and sorted in the 2D class averages. The initial model was built using EMAN2 (19). A total of 10,282 particles in 11 classes were used for 3D refinement by imposing C1 symmetry. The Gold-standard FSC was calculated using the two halve maps independently reconstructed from the particle images (Supplementary Fig. S2). Amplitude correction was performed without masking the structure. The obtained structure was rendered with UCSF Chimera (20).

### Dissociation analyses of TANAV particles

Purified TANAV particles were incubated at different pH levels as follows: pH 8, 10 mM Tris-HCl, 100 mM NaCl, 1 mM EDTA; pH 6, 250 mM sodium acetate, 10 mM Tris-HCl, 100 mM NaCl, 1 mM EDTA; pH 5, 250 mM sodium acetate, 10 mM Tris-HCl, 100 mM NaCl, 1 mM EDTA; pH ∼3, 50 mM HCl; pH ∼11 50 mM NaOH. Incubation parameters also included detergent concentrations ranging from 0.01 to 0.5% NP-40, and temperatures at 37°C or 50°C for 2 or 18 hr. After treatment, the virus was loaded onto a grid and negatively stained. Negatively stained virus particles were observed on a 30 kV scanning electron microscope (Thermo Fisher Scientific, Quanta FEG 650) in scanning transmission electron microscopy (STEM) mode. All images were taken at a nominal magnification of 300,000 ×.

### Cryo-electron tomography of bullet-like TANAV particles

Purified TANAV particles were incubated overnight at 37°C in pH 5, 250 mM sodium acetate, 10 mM Tris-HCl, 100 mM NaCl, and 1 mM EDTA. The incubated sample was loaded onto a holey carbon grid and flash-frozen for collecting tilt series of the bullet-like TANAV particles by aforementioned cryo-EM equipment. The tilt series (±60°, 2° angular increments) of the bullet-like TANAV particles images were collected at a nominal magnification of 20,000 × (2.82 Å/pixel on specimen) using a low dose mode, where the total electron dose was less than 100 e^-^/Å^-2^ on the specimen. Image alignment and tomographic reconstruction were performed by IMOD software (21) using fiducial markers. The sub-tomogram averaging of 104 sub-volumes was performed using RELION 2.0 (18, 22). After iterative 2D and 3D classifications and map refinement, 27 sub-volumes were selected and used for the final 3D averaged map.

### Asymmetric and helical reconstruction of tubular TANAV particles

Purified TANAV particles were incubated at 50°C for 2 hr in pH 5, 0.01% NP-40, 250 mM sodium acetate, 10 mM Tris-HCl, 100 mM NaCl, and 1 mM EDTA,. The incubated sample was loaded onto a holey carbon grid and flash-frozen for collecting cryo-EM images of the tubular TANAV particles. The symmetric (C1) and helical reconstruction were performed using the boxed regions in Fig. 5B using RELION 3.0 (23). Helical parameters were estimated from a polar diagram (surface pattern) of the tubular structure.

## Results

### Cryo-EM structure of the TANAV particles

We first observed purified TANAV particles in near native conditions using cryo-EM (Fig. 1A). In a vitreous thin-ice film, the particles displayed round or elliptical shapes, representing standing (top view) and laid-down (side view) particles, respectively (Fig. 1A). The top view of the particles showed a bumpy outline that appeared to be consistent with a certain periodicity (Fig. 1B, top view). In the side view, the particles formed a regular elliptical shape (Figs. 1B, side view), though they appeared deformed when viewed using negative stain EM (3). Using cryo-EM, a short extension from one end of the elliptical particle can be visualized (Fig. 1B, yellow arrows in side view). This structure corresponds to the short projection that was previously observed in negatively stained TANAV particles (3). 2D class-averaged particles are shown in Figure 1C. The class-averaged images illustrate apparent top and side views in addition to those from intermediate angles (Fig. 1C). Periodic bumps, 29 in all, can be seen in the top view (Fig. 1Da, yellow), forming a round shape with a maximum diameter of 37 nm (Fig. 1Da). In the side view, the short projection is visible on one vertex of the 55-nm-long elliptical core (Fig. 1Db). The smeared short projection in the 2D average appears to be branched and tufted (Fig. 1Db, white arrow).

**Fig. 1.**
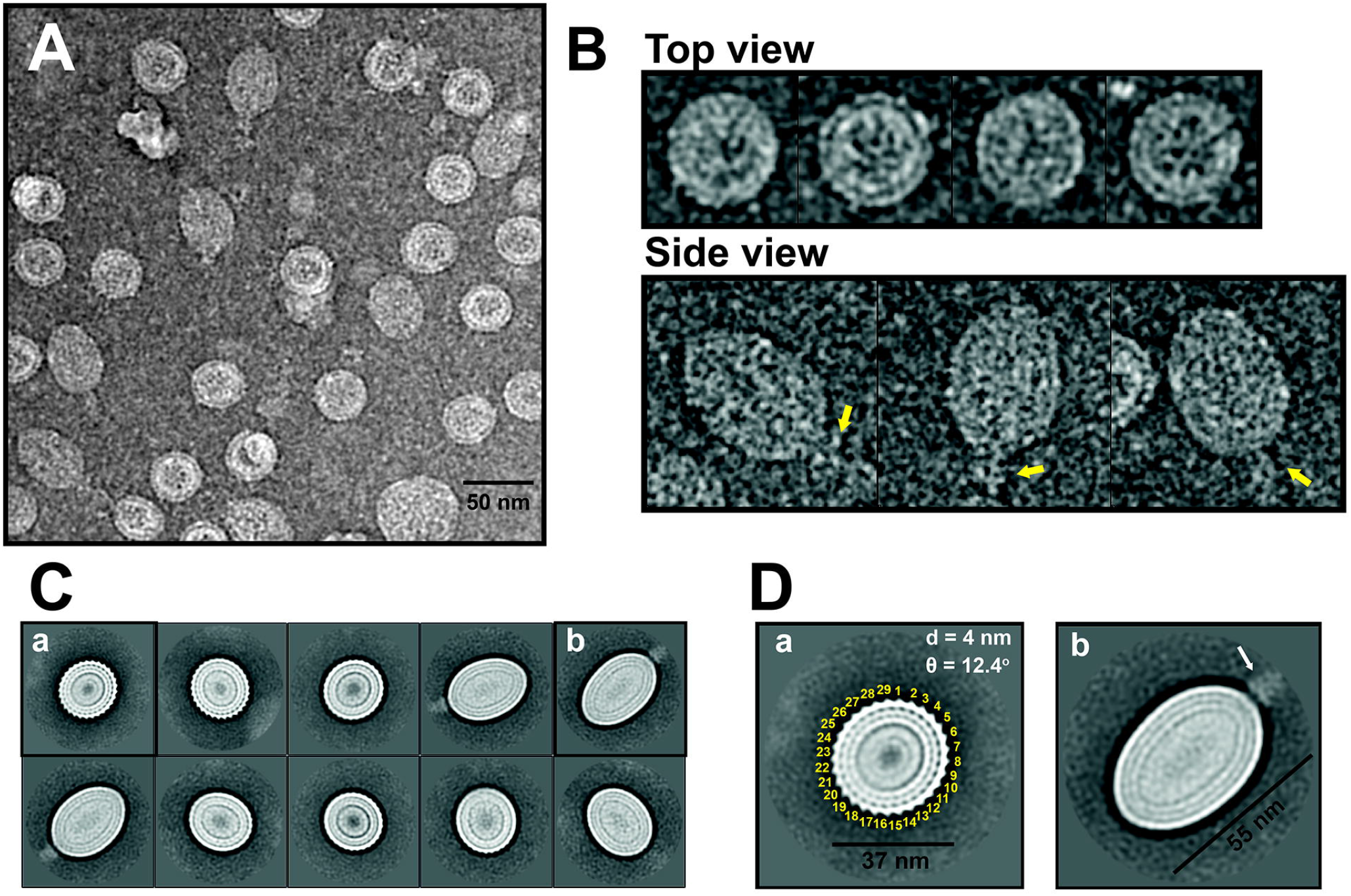
Cryo-EM images and 2D-class averaging of the TANAV particles. A) Raw image of TANAV particles under cryogenic conditions. The image contrast is reversed. B) Representative standing (top view) and laid down (side view) TANAV particles. Yellow arrows indicate the short projection on the particles. C) 2D class-averaged particle images. D) Close-up view of standing (a) and laid down (b) 2D class-averaged particle images in (C). Assigned numbers of periodical bumps are labeled. Distance and angle between adjacent bumps are included in (a). White arrow indicates the short projection in (b).

The subtle structural heterogeneity of enveloped TANAV particles posed a challenge for reconstruction in 3D using cryo-EM SPA. Since it was unclear whether elliptical TANAV particles have symmetry, asymmetrical reconstruction (C1) was initially performed to derive the entire 3D structure of the TANAV particle. The resolution was estimated to be 23.6 Å without imposing symmetry (C1) by using a gold-standard Fourier shell correlation (FSC) cutoff of 0.143 (Supplementary Fig. S2). The C1 reconstruction showed an elliptical core and a short projection (Fig. 2). A center slice along the short axis of the elliptical core exhibited regular bumps as seen in the 2D averaged classes of TANAV particles (Figs. 1D, 2E and 2G). The number of bumps at the middle of the elliptical core is confirmed to be 29 by assuming optimal rotational symmetry (Fig. 2G, Supplementary Fig. S3). However, the number of the bumps at the distal parts of the elliptical core gradually reduce to 5 (Supplementary Fig. S2). These results suggest that the capsid of the elliptical core has a spiral structure. Further, the base of the short projection seems to show C3 symmetry, while the distal edge shows roughly C2 symmetry (Supplementary Fig. S3). To enhance the structural features of the TANAV particle, different rotational symmetries (C29: center – C2: projection) were temporally imposed on each slice of the C1 reconstruction across the major axis of the TANAV particle (Figs. 2C, D, G, H, Supplementary Fig. S3). TANAV particles are composed of three layers in the periphery of the particle (1-3 in Fig. 2G) and of one interior fragile core representing viral RNA (Figs 2E-H). The outermost layer is thicker and clearer than the other two peripheral layers (Figs. 2E-H). The center of the interior core does not show an intensity, for the viral RNA seems to be absent or loosely packed (Figs. 2E-H). The layer structures near the major axis of the elliptic core look slightly different from the others, in that the second layer is disconnected near the short projection while it has more density at the opposite edge (Red brackets in Fig. 2H).

**Fig. 2.**
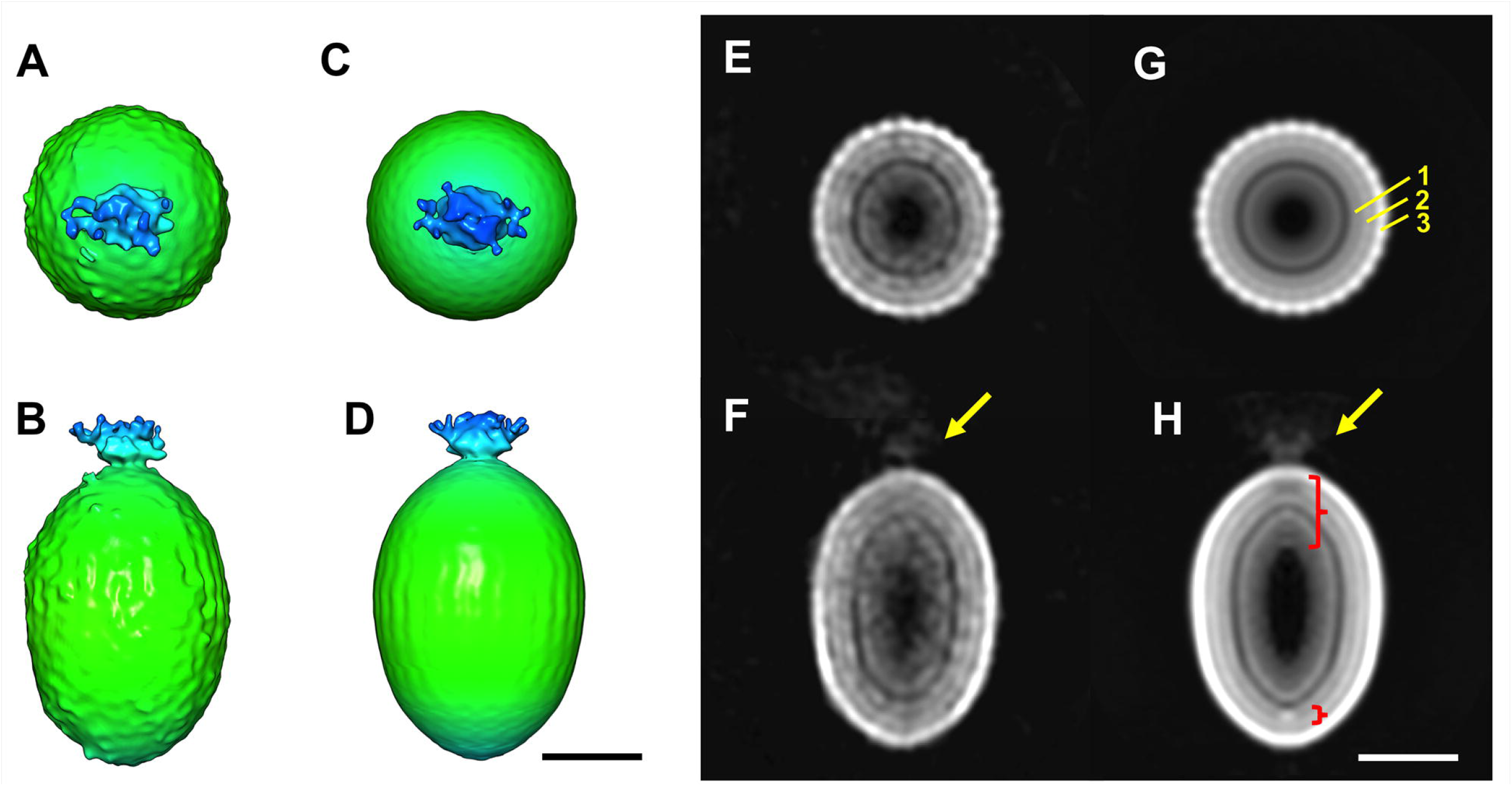
Single Particle Reconstruction of the TANAV particle. A, B) Non-symmetrized C1 reconstruction. C, D) Symmetrized model of the C1 reconstruction. To enhance the structural features, putative rotational symmetries were applied for each Z slices. Z-axis is the major axis of the oval TANAV particles. E, F) Center slice of (A, B). G, H) Center slice of (C, D). Yellow arrows indicate the short projection. Three outermost layers are shown in G). The partial modification of the layers are shown in H) (red blankets). The scale bar is 20 nm. The contour level is set to 1 σ.

### Structural dissociation analyses of the TANAV particles

TANAV was exposed to various pH levels, detergent concentrations, and temperatures to dissociate the structure of the TANAV particle and elucidate the geometrical characteristics of its elliptical envelope. Elliptical TANAV particles were shown to be stable under thermal conditions, remaining intact at 50°C (Supplementary Fig. S4A). Different concentrations (0.01%-0.5%) of NP-40 detergent were also used to dissociate the enveloped particles. TANAV particles did not dissociate at 0.01% NP-40, but they did degrade at concentrations above 0.05% NP-40 (Supplementary Fig. S4B). The most significant structural dissociations were observed when the buffer pH was set to acidic (Fig. 3A). Intact TANAV particles were originally suspended in a pH 8 buffer (Fig. 3A, left). Elliptical structures were maintained down to pH 6 (Fig. 3A, middle), but they transformed into a bullet-like shape at pH 5 (Fig. 3A, right, yellow arrows). These bullet-like TANAV particles did not include the short projection that was observed at pH 8 and pH 6 (Fig. 3A), suggesting that it was removed at pH 5. TANAV particles can be degraded under both extremely acidic (50 mM HCl, pH = ∼3) and extremely basic (50 mM NaOH, pH = ∼11) conditions, though certain particle structures are retained even at an extremely low pH (Supplementary Fig. S4C). Further dissociation analysis was performed under acidic conditions containing 0.01% NP-40 (Fig. 3B). At mid-range pH values, 0.01% NP-40 did not affect the structure of TANAV particles. However, particles did tend to aggregate at pH 6 (Fig. 3B, middle), and at pH 5 there was significant particle dissociation along with the formation of tubular structures (Fig. 3B, right). Similar to the bullet-like structures observed at pH 5 (Fig. 3A, right), these structures failed to show the short projection observed at pH 6 and pH 8 (Fig. 3B, right).

**Fig. 3.**
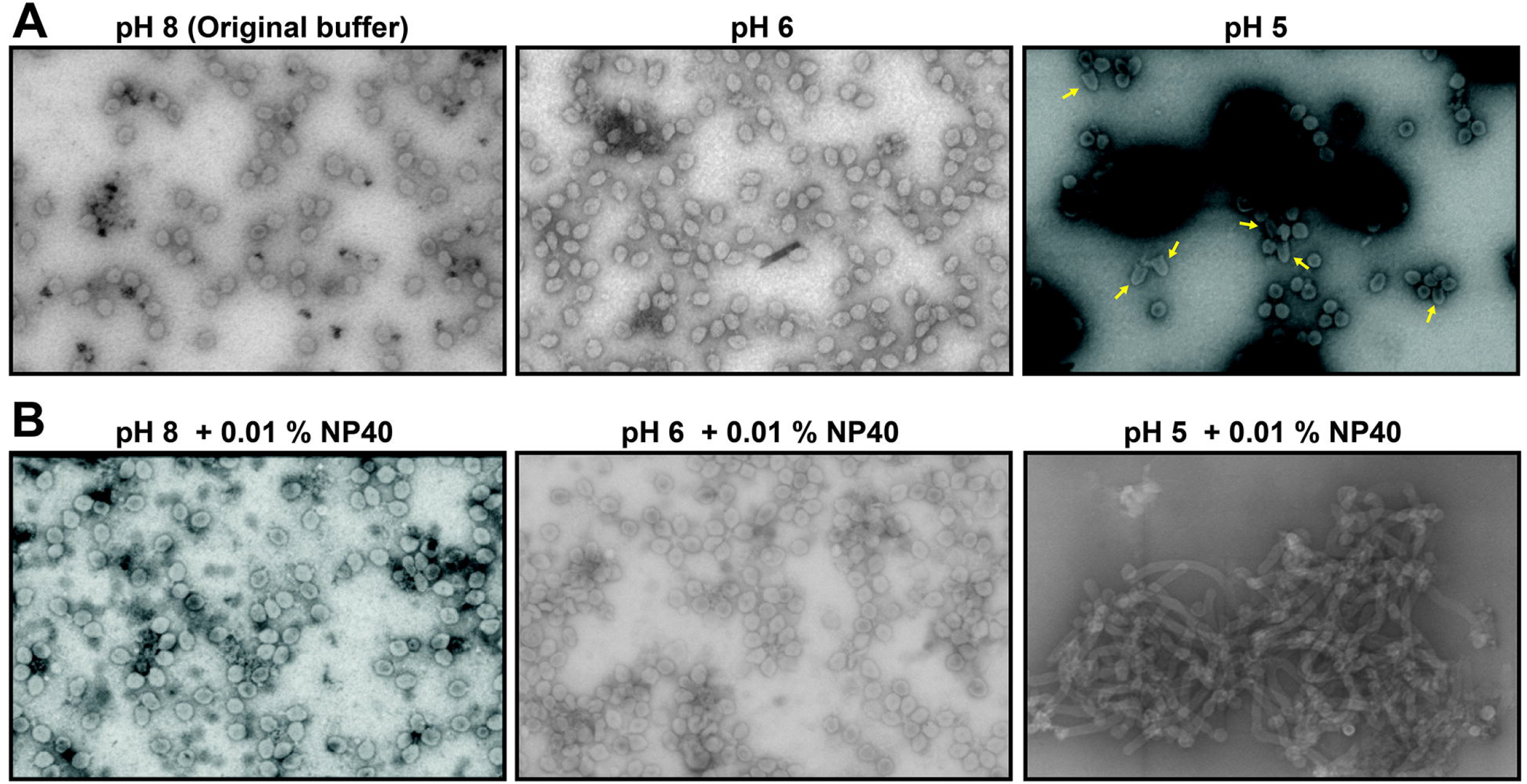
TANAV particles under acidic and low detergent conditions. STEM images (300,000 ×) of negatively stained TANAV particles are shown. A) pH 8, pH 6, and pH 5 buffers at 37°C for 18 hr. B) 0.01% NP-40 is further added to pH 8, pH 6, and pH 5 buffers at 50°C for 2 hr. Bullet-like particles are indicated with yellow arrows in the pH 5 panel (A, right). Tubular TANAV particles appear at pH 5 + 0.01% NP-40 (B, right).

### Cryo-EM structures of the transformed TANAV particles

To investigate 3D structures of bullet-like and tubular TANAV particles, their raw particle images were collected at a near-native condition using cryo-EM (Figs. 4, 5).

**Fig. 4.**
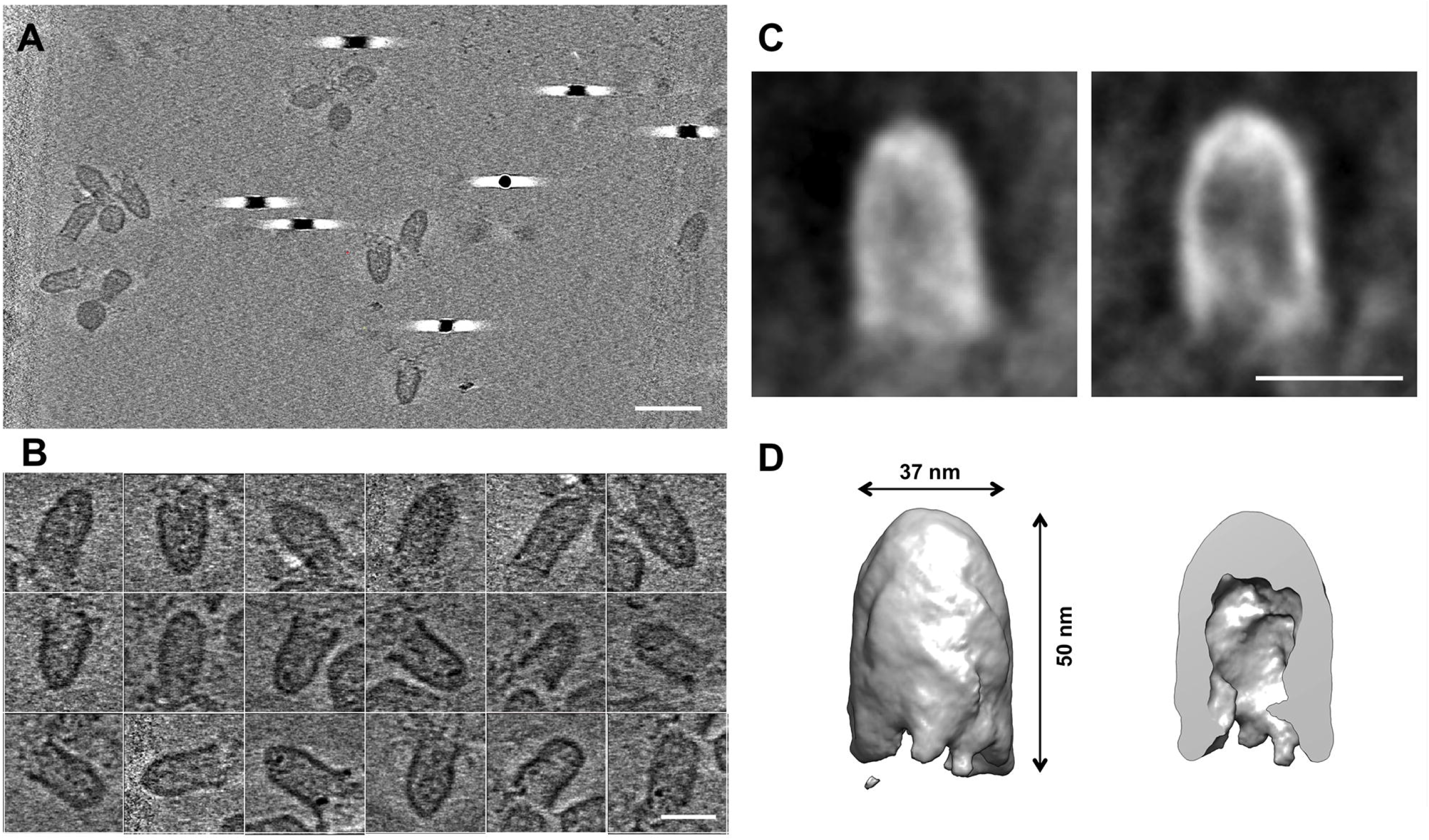
Cryo-ET of the bullet-like TANAV particles. The TANAV particles were incubated in pH 5 buffer at 37°C overnight for dissociating to form the bullet-like particles. A) Reconstructed cryo-EM tomogram of the bullet-like TANAV particles. The scale bar is 100 nm. B) Cryo-EM raw images of the bullet-like TANAV particles. C) Sub-tomogram averaged structure of the bullet-like TANAV particle (left) and its cross-section (right). The scale bar is 40 nm in (B, C). D) Isosurface view of the bullet-like TANAV (left) and its cross-section (right).

**Fig. 5.**
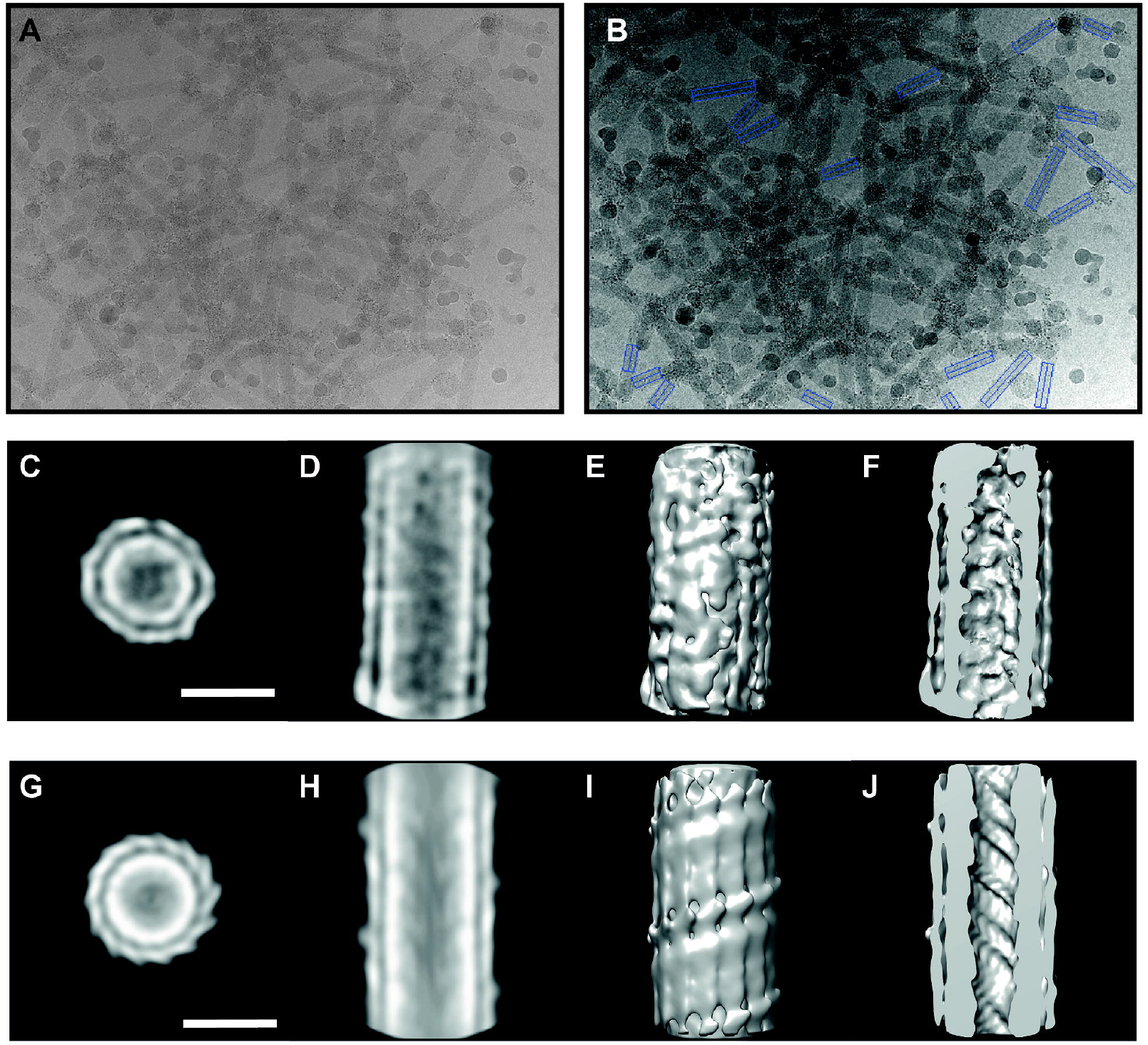
Single particle analysis of the tubular TANAV particles. A, B) Cryo-EM raw images. Blue boxes in B) show the regions used for the single particle analysis. C-J) C1 (C-F) and helical (G-J) reconstructions of the tubular TANAV particle. The 2D class-averaged projections of the C1 and helical reconstructions are shown from top view in (C, G) and side view in (D, H). The 3D cryo-EM structure of the C1 and helical reconstructions of the tubular TANAV particle are shown in (E, I) and their cross-sections are in (F, J). The scale bars are 20 nm.

The bullet-like TANAV particles generated at pH 5 were not homogenous enough to determine their structure using cryo-EM SPA (Figs. 4A, 4B), and thus cryo-ET and subtomogram averaging were applied to enable structural determination in 3D (Figs. 4C, 4D). The length and width of the bullet-like TANAV particles correspond well to the major and minor axes of the original elliptical particles (Figs. 1D and 4D). However, the interior density of the bullet-like particles was low compared to that of the elliptical particles (Figs. 2F, 2H and 4C). The short projection of the elliptical TANAV particles also disappeared in the bullet-like particles (Fig. 4). These observations suggest that the structure of the bullet-like particles after releasing the viral RNA involves removal of the short projection and opening of the elliptical core. The bullet-like TANAV particles showed only one thick layer, suggesting that the structured surface layer of the elliptical TANAV particles was disordered (Figs. 2E-H, 4B-D).

It was, on the other hand, possible to use cryo-EM SPA to reconstruct the tubular TANAV particles generated at pH 5 with 0.01% NP-40 by imposing C1 symmetry or helical symmetry (Fig. 5). The tube diameter of approximately 25 nm was much smaller than that of the elliptical and bullet-like TANAV particles, each of which had a diameter of approximately 37 nm (Figs. 2B, 2D, 4D, 5C-J). The fine structure of tubular TANAV particles was enhanced by imposing helical symmetry, and the number of bumps observed on the surfaces of the tubular TANAV particles was almost half (14 bumps) the number detected at the centers of the elliptical TANAV particles (29 bumps) (Figs. 2E, 5G). Tubular TANAV particles showed only two layers: an outer thin layer and an interior thick layer (Figs. 5D, F, H, J). The middle of the tube was empty.

## Discussion

Our study reveals, for the first time, the 3D structure of elliptical and enveloped TANAV particles (Figs. 1, 2). At high resolution, particle heterogeneity hinders the determination of 3D structure, though it is still possible to reveal the novel geometry of elliptical TANAV particles as well as their transformations. Proposed structural models of intact ellipsoid, transformed bullet-like, and tubular TANAV particles are summarized in Fig. 6. In the top views of the 2D class-averaged and 3D reconstructed images, periodic bumps appear on the surface of the elliptical core (Figs. 1C-D, 2E, 2G), implying that envelope proteins are arranged with a spiral or cylindrical symmetry through some or all of the particle’s elliptical core. TANAV has been reported to express an ORF3-encoding single structural protein that forms the elliptical enveloped core (3,8,13) (Supplementary Fig. S5). The 3D reconstruction of TANAV particles indicates the presence of distinct rotational symmetries along the major axes of elliptical particles (Supplementary Fig. S3). The rearrangement of TANAV structural proteins in tubular particles appears to follow helical symmetry (Figs. 5G-J), suggesting that the ORF3 protein has the ability to align helically. Based upon these results, we assume that the periodic bumps correspond to the helical arrangement of structural proteins and that the number of the bumps/structural proteins reduces at each end of the elliptical core due to this geometry. In other words, it exhibits a spiral array of structural proteins along the surface of the viral envelope. Bullet-like particles are hatched on one side of the elliptical core through removal of the short projection and release of the spiral geometry. Tubular particles can squeeze the inherent helical array of 29 structural proteins per turn, resulting in 14 structural proteins per turn. Further studies are needed to understand the mechanisms of these dynamic transformations.

**Fig. 6.**
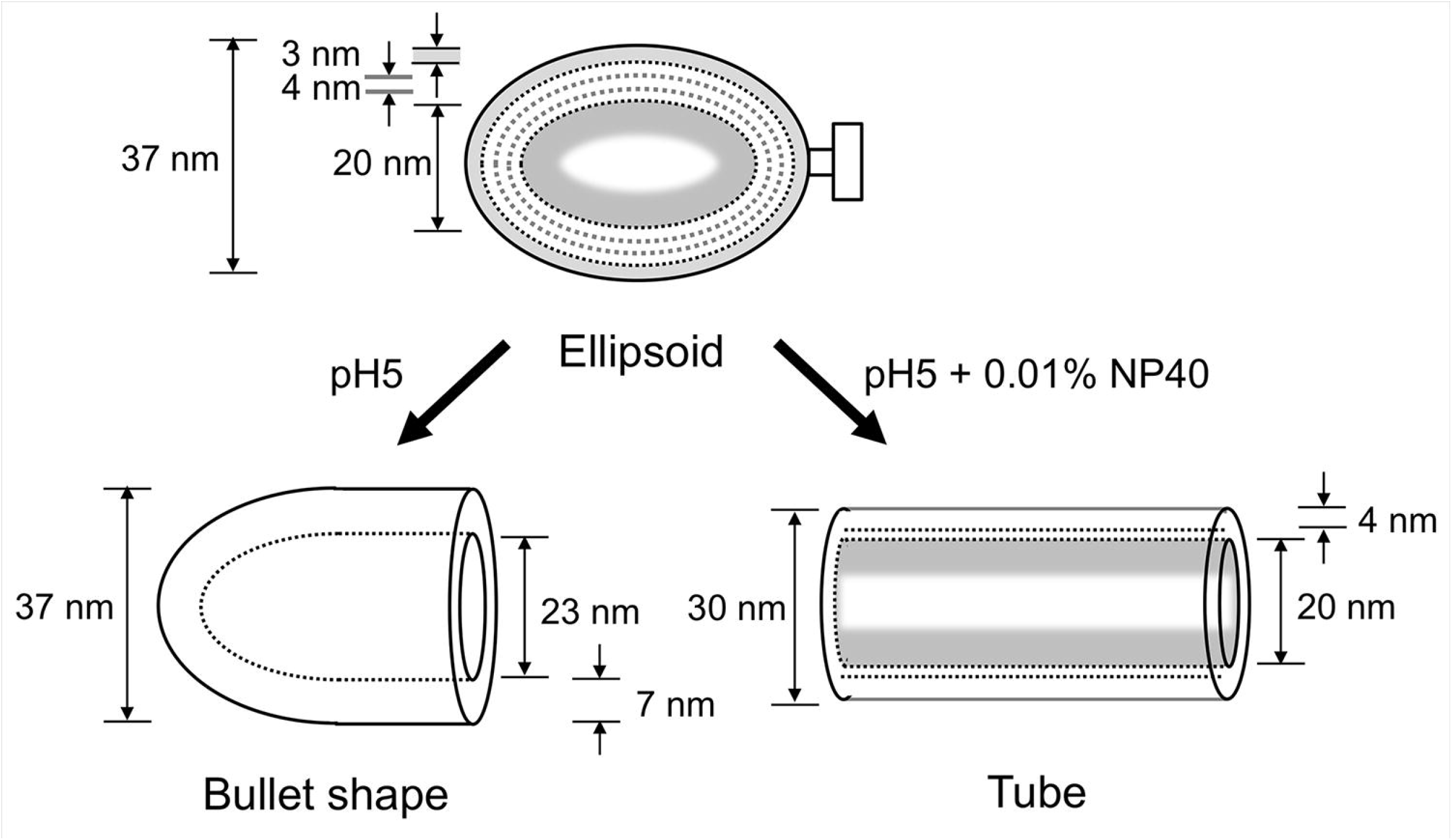
Schematic models of the intact and transformed TANAV particles. The bullet-like and tubular particles are formed at pH 5 without or with 0.01% NP-40, respectively. The ellipsoid particle (37 nm long) consists of the outer three layers (the outermost thick layer of 3 nm in width and the thin 2nd and 3rd layers of 4 nm in total width) and the one interior fragile layer. The bullet-like particle (37 nm long) consists of one thick layer (7 nm in width). The interior fragile layer of the ellipsoid particle is absent in the interior space of the bullet-like particle. The tubular particle (30 nm in diameter) consists of the outermost layer (4 nm in width) and the thicker interior layer.

The layered structure of the TANAV envelope is dynamically changed during structural dissociation (Fig. 6). As revealed by hydropathy plot, the ORF3-encoding structural protein is a transmembrane protein with large ectodomains (Supplementary Fig. 5A and B). The three peripheral layers of the elliptical TANAV particle are thus assumed to be ORF3 proteins and a lipid bilayer. According to a topology model based on hydropathy analysis (Supplementary Fig. S5C), the outermost thick layer is composed of the ectodomains of ORF3 proteins whereas the two inner layers are transmembrane domains that are embedded in the lipid bilayer of the viral envelope (Fig. 6). The distance between the two inner layers (4 nm) is a regular width of the lipid bilayer. Interestingly, the total width (3 + 4 nm) of the outermost layer and the two inner layers of the elliptical particles was similar to the width of the disordered thick layer (7 nm) of the bullet-like particle, despite the presence or absence of viral RNA (Fig. 6). Considering the structural reorganization of ORF3 proteins in bullet-like particles, the short projection of the elliptical TANAV particles not only order the ORF3 proteins but also play an important role in packaging the viral genome into the elliptical core. In contrast, tubular particles appear to have two layers (Figs. 5C-J, 6). The dynamic structural rearrangement of ORF3 proteins under the low-detergent condition reveals the different number of the layers in tubular particles. Map intensity and helical geometry tentatively suggests that the outer layer (4 nm in width) is composed of a part of the helically arranged ORF3 protein (Fig. 6), while the thick interior layer (20 nm diameter in total) could consist of another part of the ORF3 protein along with viral RNA. Further studies are, however, needed to refine the detailed structural model.

Certain archaeal viruses display a lemon-shaped or helical capped structure, such as His1 (*Haloarcula hispanica*virus 1) and APBV-1 (*Aeropyrum pernix* bacilliform virus 1), respectively (24, 25). The pointed cap structure of APBV-1 was resolved at 8 Å resolution, suggesting that the long helical core structure assembled by similar VP1 proteins was built by a similar mechanism (25). His1 particles change from a lemon-shaped overall core to a tubular shape under high temperature (80°C) and low concentrations of detergent (24), implying a potential structural similarity to insect TANAV particles. A remarkable difference, however, is that the tubular formation of TANAV occurs regardless of temperature (Fig. 3, Supplementary Fig. S4). The insect virus TANAV may have been adapted to the relatively low body temperature of mosquitoes, whereas the thermophilic archaea virus His 1, which shows a temperature-dependent structural change, is likely the result of adaptation to a heated host environment. Tubular structures were reported in other negeviruses found in the infected cells (10), but tubular TANAV particles have not been observed in TANAV-infected mosquito cells. Such particles instead exhibiting a round or oval shape imply that they are intact TANAV particles in the host cell (Supplementary Fig. S6). The physiological function of the tubular structure in TANAV is not clear at present, but it may be a remnant of negevirus assembly. Although it would be interesting to perform an in-depth characterization with respect to the protein and genome composition of these TANAV conformational variants, at present, it has not been successful in purifying these conformational variants due to particle aggregation under low pH and/or detergent conditions.

Many enveloped viruses, including mosquito viruses, alter their structural conformation during intracellular endosomal acidification for cell entry (26). In the case of TANAV, the short projection dissociates under acidic conditions to yield a bullet-like shape and release the viral RNA (Fig. 3A, right). The glycoprotein of negeviruses forming the short projection might have evolved to facilitate infection of insect cells, because the short projection is not identified in phylogenetically related plant-specific viruses (14, 15). Unlike such plant-specific viruses, TANAV particles may have used to have a short projection to acquired sensitivity to pH for membrane fusion or genome release during endosomal uptake. If negeviruses derived from plant viruses adapted to mosquito hosts by acquiring this short projection, some negaviruses could infect plants, as do other plant (+) ssRNA viruses. Negeviruses are currently thought to infect only mosquitoes and other arthropods, but further studies are necessary to clarify their ability to infect certain plants.

## Acknowledgements

This work was supported by the following agencies: Vetenskapsrådet (VR)/The Swedish Research Council (to K.O., grant number: 2018-03387), the Swedish Foundation for International Cooperation in Research and Higher Education (STINT) (to Janos Hajdu and K.O., grant number: JA2014-5721), FORMAS research grant from the Swedish Research Council for Environment, Agricultural Sciences and Spatial Planning (to K.O., grant number: 2018-00421), Royal Swedish Academy of Sciences (to K.O., grant number: BS2018-0053), KAKENHI from the Ministry of Education, Culture, Sports, Science and Technology of Japan (to N.M., grant Number: 25251009) and the Collaborative Study Program of National Institute for Physiological Sciences (to K.O., grant Number: 2016-No. 38).

## Author Contributions Statement

K.O., and S.I. prepared samples. K.O., C.S. and K.M. wrote the manuscript. K.O., and K.M designed the experiments. K.O, C.S., N.M., and K.M. collected the cryo-EM data and analysed the data. C.S., M.S., N.M., T.N., K.M., and S.I. provided technical support on the experiments. All of the authors discussed the results and proofread on the manuscript.

## Accession codes

The 3D cryo-EM map of the TANAV has been deposited in the Electron Microscopy Data Bank under the accession number EMD-10601.

