## Supplementary figures for "Structure and its transformation of elliptical nege-like virus Tanay virus"

**Supplementary information**

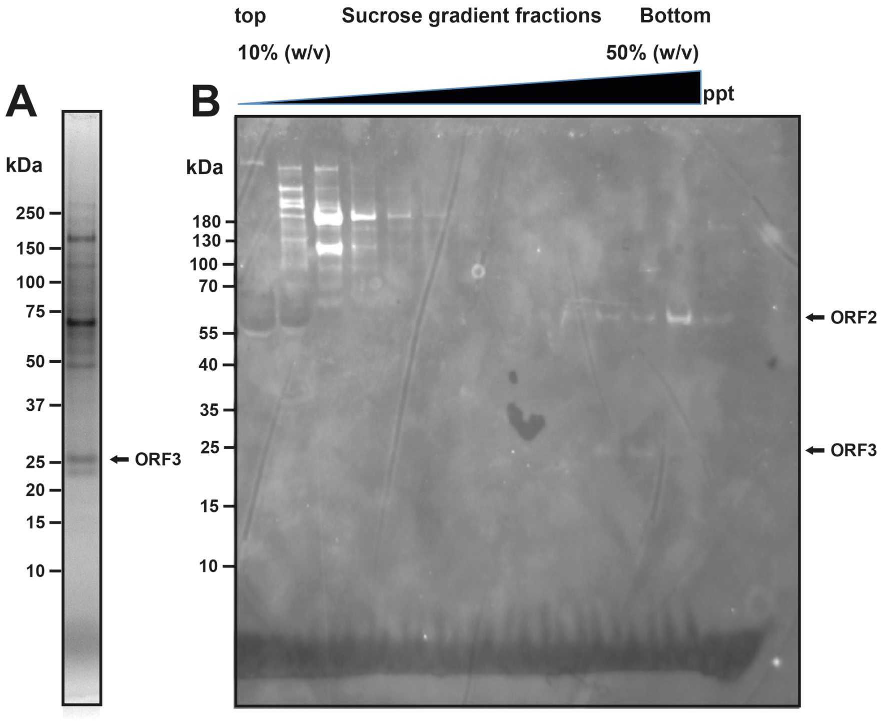


**Supplementary Figure S1. Purified TANAV sample.** A) Coomassie-stained protein bands in the purified TANAV sample. B) Lectin blotting of the sucrose gradient fractions for the TANAV purification. For the lectin blotting, the SDS-PAGE gel bands of the sucrose gradient fractions were transferred to nitrocellulose membrane in transfer buffer (25 mM Tris base, 192 mM Glycine, 10% (v/v) Methanol) using Bio-rad turbo blot. The membrane was blocked for 2 hr in 3% (w/v) skimmed milk at RT. The membrane was treated with a lectin blotting solution (100 µg/mL Alexa488-conjugated concavalin A, 0.3% (w/v) skimmed milk in PBST) for 2 hr at room temperature (RT). After rinsed 3 timed in PBST, the membrane was imaged using BioRad ChemiDoc to detect Alexa488 fluorescent signals.

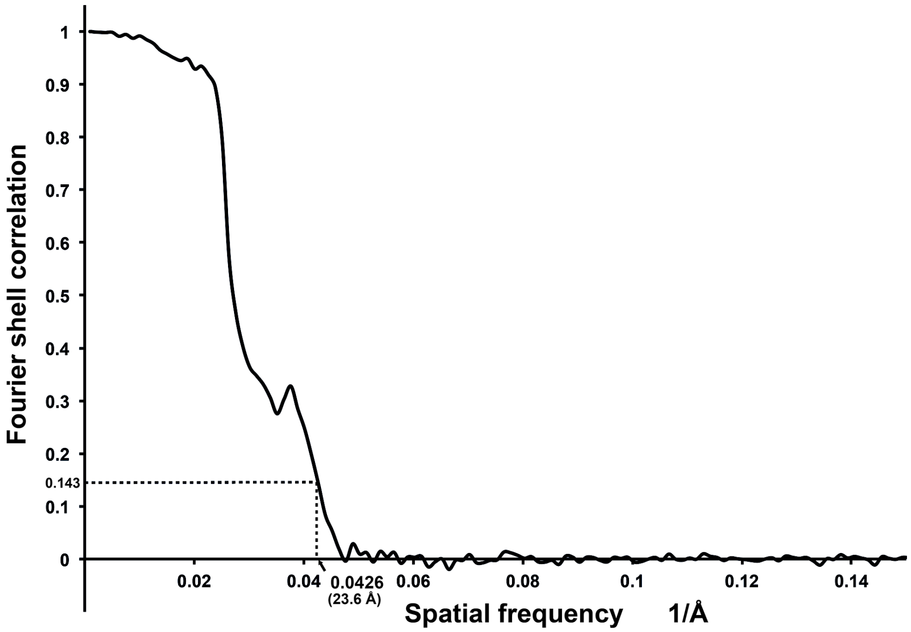


**Supplementary Figure S2. Estimated resolution of reconstructed cryo-EM structure of the TANAV particle.** The resolution of the C1 reconstruction was calculated to be 23.6 Å using 0.143 cutoff of gold-standard Fourier shell correlation.

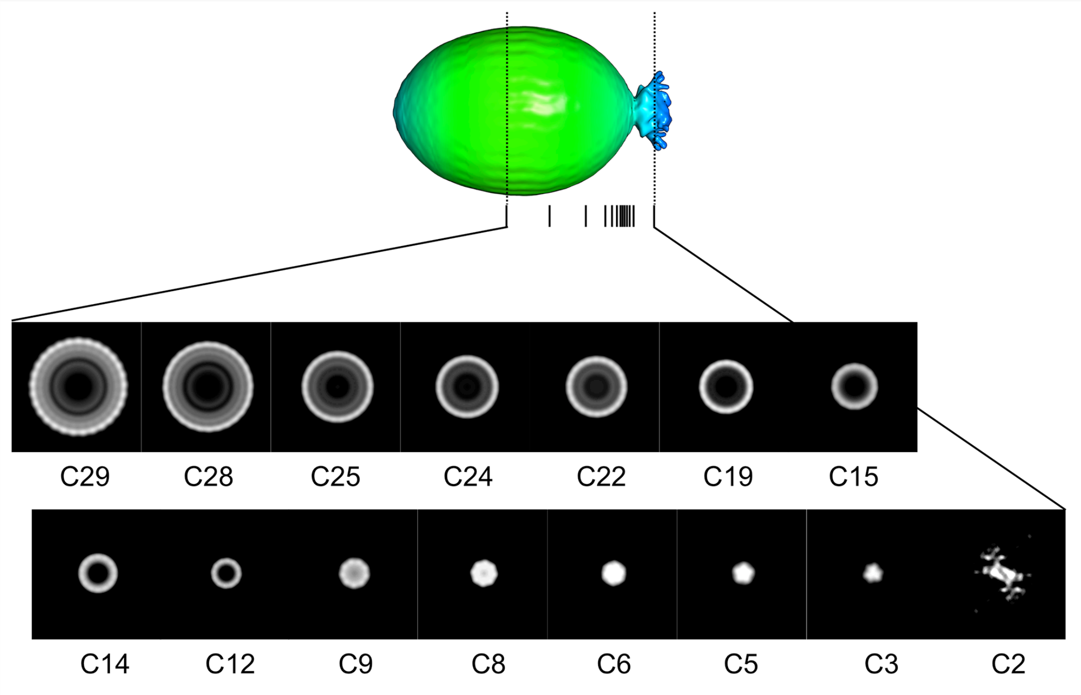


**Supplementary Figure S3. Symmetry operation to the C1 reconstructed model of the TANAV particle.** Putative rotational symmetries (C29 – C2) were independently applied to different z-axis positions from the middle to the edge of the elliptic core. Each of the cross-sections that are imposed different rotational symmetry is displayed. The imposed C29-C5 rotational symmetries in the elliptic fore of the TANAV particle enhanced the structure features from the middle to the edge. The result suggests that the elliptic core of the TANAV particle form a spiral structure. Furthermore, the base of the short projection seems to show a C3 symmetry, while the distal edge is roughly a C2 symmetry.

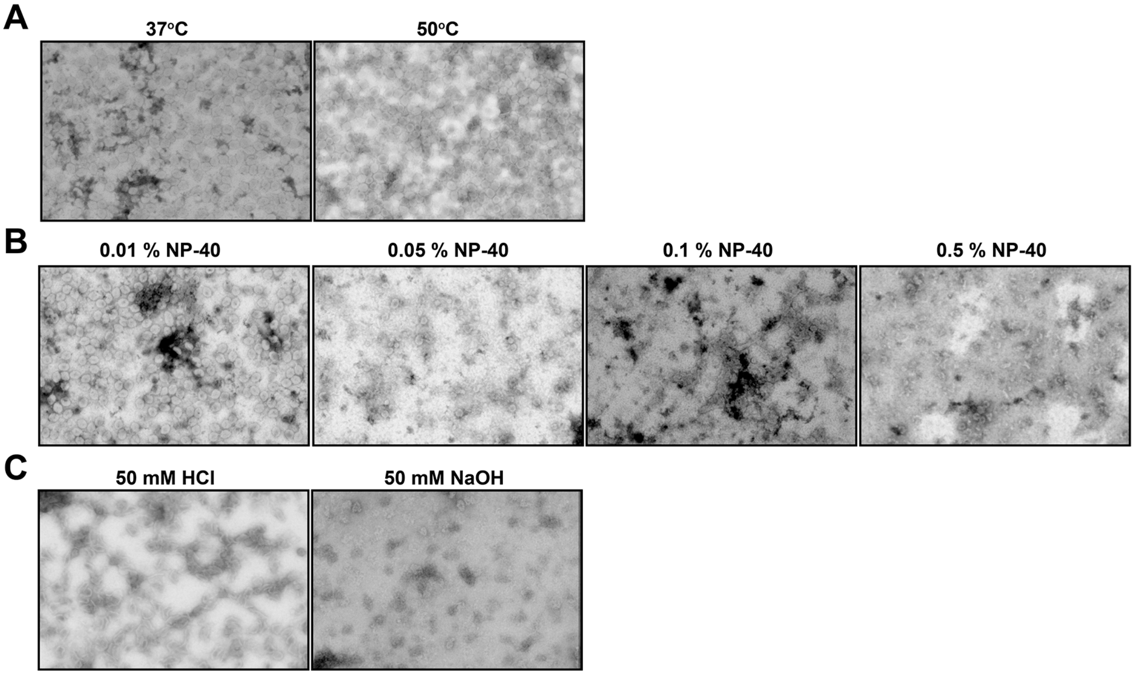


**Supplementary Figure S4. TANAV particles under heating, various detergent conditions, and extremely basic or acidic pH.** STEM images (300,000 x) of negatively stained TANAV particles are shown. A) 37°C (left) and 50°C (right) at pH 8 for 2 hr. B) 0.01 – 0.5% NP-40 at 37°C, pH8 for 2 hr. C) 50 mM HCl (pH ~3) (left) and 50 mM NaOH (pH ~11) (right) at 37°C for 2 hr.

**A**

```
                                TM helix 1
1   MVLKRTTVAA RKVTSSKTAG RPVKVTEIRV TPKKTATKTS GKTERKSKDF 50
51  DFSAYFDYTV KVLTNPSYVI FFVCAAYLCF NYMHEHSNSH LWTFVQNFVN 100
                         TM helix 2
101 KFTTFKTSAC SILNMMLCFV PFIPAIVSVP SKNRSLTIIC TIAYYMFIPE 150
                                              TM helix 3
151 RTPYEYLAHG IIVFLILKTN NQQYRMIGVA LLFLTYIMQF VIPLPMPDVS 200
201 YVCNGTSLVA APKSQ
```

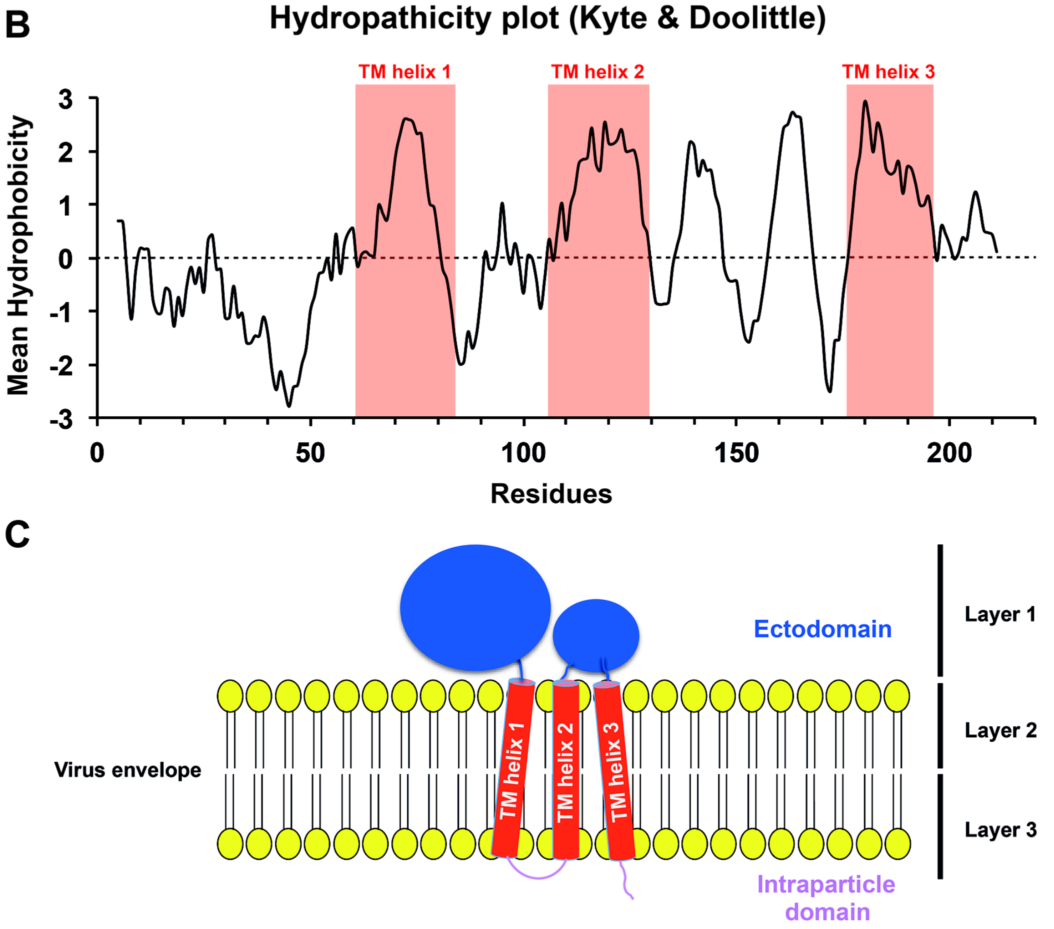


**Supplementary Figure S5. Hydropathy analysis and predicted topology of the TANAV major envelope protein.** A) A prediction of transmembrane helices (TM helices) in an amino acid sequence of TANAV ORF3 major envelope protein. Three TM helices are predicted in the amino acid sequence using TMHMM Server v.2.0 (Red boxes) (1). The expected sequences of ectodomain and intraparticle domain are colored in blue and pink, respectively. B) Hydropathy plot of the TANAV ORF3 protein. The predicted three TM helices correspond to the red regions. C) A topology model of the TANAV envelope protein. The colors of the model correspond to those in (A). The three outer layers observed in Figure 1C and 1D are assumed to correspond to the labeled layer 1-3 of the aligned TANAV major envelope ORF3 proteins in the virus envelope.

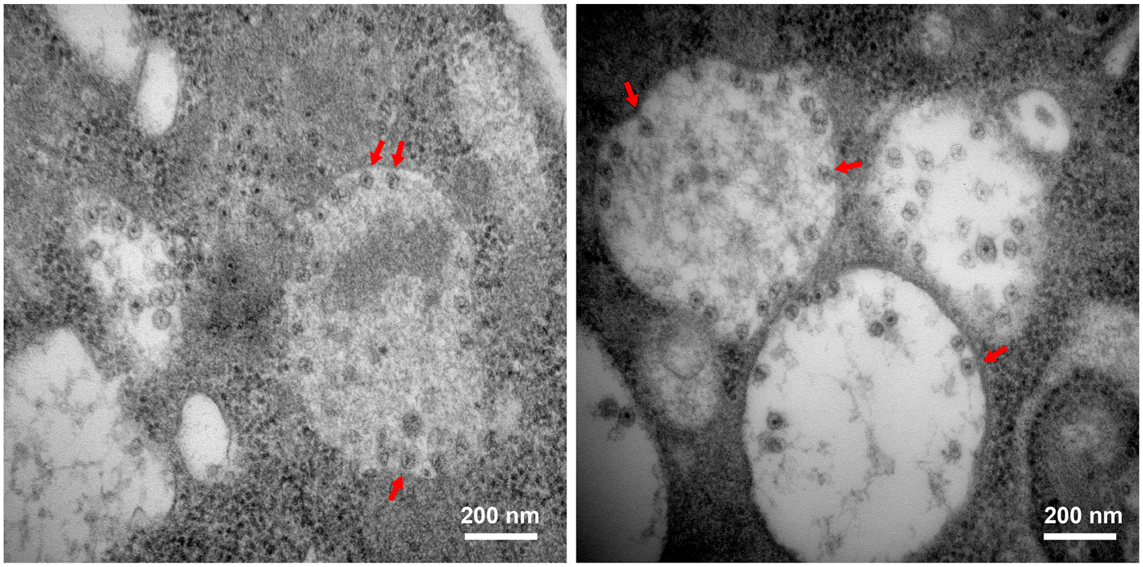


**Supplementary Figure S6. Virus factories of the TANAV particles in TANAV-infected C6/36 cells.** The purified TANAV particles were inoculated to C6/36 cells and incubated at 28°C and 5% $CO_2$ for 24 hr. The TANAV-infected cells were washed by PBS (-), and then fixed with 2% (v/v) glutaraldehyde and 2% (v/v) paraformaldehyde in PBS (-) for 2 hr. The fixed cells were washed 6 times by 1 mL of 0.1 M sucrose in 0.1 M phosphate buffer. The cells were treated by osmium solution and embedded in resin for preparing thin sections of the TANAV-infected cells. The ultra-thin sections were observed in a transmission electron microscope (100,000 ×, JEM-1010). The assembling TANAV particles in the virus factories are shown in red arrows.
